# Dapagliflozin improves endothelial integrity and hemodynamics in endotoxin treated mice through an apolipoprotein M dependent pathway

**DOI:** 10.1101/2022.04.27.489709

**Authors:** Carla Valenzuela Ripoll, Zhen Guo, Tripti Kumari, Kana N. Miyata, Mualla Ozcan, Ahmed Diab, Amanda Girardi, Li He, Attila Kovacs, Carla Weinheimer, Jess Nigro, Jan Oscarsson, Russell Esterline, Joel Schilling, Mikhail Kosiborod, Christina Christoffersen, Jaehyung Cho, Ali Javaheri

## Abstract

**Rationale:** Sodium-glucose co-transporter inhibitors (SGLT2i) are under active clinical investigation in patients with acute inflammatory conditions, based on their clinical cardio-and nephroprotective effects, and a pre-clinical study that demonstrated SGLT2i improve renal outcomes and survival in a lipopolysaccharide (LPS) model. However, a unified mechanism that explains how SGLT2i could prevent hemodynamic consequences of inflammatory conditions has not been described. Apolipoprotein M (ApoM) is inversely associated with mortality in inflammatory conditions and improves cardiac function in endotoxin-treated mice via sphingosine-1-phosphate (S1P) signaling.

**Objective:** Test the hypothesis that pre-treatment with SGLT2i dapagliflozin (Dapa) improves hemodynamics in endotoxin-treated mice via the ApoM/S1P pathway.

**Methods and Results:** Mice with diet-induced obesity were gavaged with vehicle or Dapa for 4 days prior to LPS (10 mg/kg, IP). We found that mice receiving Dapa restored circulating ApoM levels, likely by increasing expression of the multi-ligand protein receptor megalin in the proximal tubules. Dapa attenuated LPS-induced reductions in cardiac dysfunction including reductions in ejection fraction, cardiac index, and coronary sinus area as well as vascular permeability as ascertained by intravital microscopy. Using both ApoM transgenic and knockout mice and S1P receptor inhibitors, we show that the ApoM/S1P pathway is important for the beneficial effects of Dapa in the LPS model.

**Conclusions:** In the setting of acute inflammation, our data suggest that SGLT2i maintains levels of megalin, leading to preservation of ApoM, which in turn promotes endothelial barrier integrity and improves hemodynamics. Our studies suggest a novel mechanism by which SGLT2i can preserve intravascular volume in the acute inflammatory setting.

## Introduction

Sodium-glucose co-transporter inhibitors (SGLT2i) such as dapagliflozin (Dapa) have well-described clinical benefits in diabetes, heart failure (HF) and chronic kidney disease (1-5). Interestingly, a recent post-hoc analysis of the Dapagliflozin in Patients with Chronic Kidney Disease trial demonstrated reduced infectious death in patients treated with SGLT2i (6), findings that are concordant with preclinical studies suggesting that SGLT2i can improve survival and renal function in a lipopolysaccharide (LPS) murine model (7). This recent post hoc analysis suggests the hypothesis that SGLT2i may reduce infectious death, but as in HF and kidney disease, the downstream mechanism by which SGLT2i leads to these improved outcomes remains enigmatic, particularly with respect to the integrative physiology.

Apolipoprotein M (ApoM) is a high-density lipoprotein-associated apolipoprotein that is secreted from hepatocytes and binds sphingosine-1-phosphate (S1P) (3, 8-14). The ApoM/SIP axis is a critical regulator of vascular inflammation (3, 8-11, 13-16), and levels of ApoM/S1P have been inversely associated with outcomes in multiple inflammatory conditions including sepsis (10, 17-19), COVID-19 (20, 21), HF (22) and type 2 diabetes (23). Given the overlap of these associations with the clinical benefits of SGLT2 inhibition, notably in type 2 diabetes and HF, we hypothesized that SGLT2 inhibition preserves ApoM levels and improves hemodynamics in the setting of acute inflammation.

## Methods

### Reagents

Dapagliflozin (Dapa) was obtained from AstraZeneca (AZ13219875-003, Stockholm, Sweden), diluted in DMSO to 100 mg/ml, and further diluted in sterile filtered water to achieve the desired concentration. Lipopolysaccharide (LPS) was purchased from Sigma Aldrich (#L2630, St. Louis, MO) and dissolved in 1X phosphate buffered saline (PBS), which was used in animal experiments. VPC23019 was purchased from Tocris (#4195, Minneapolis, MN, USA) and dissolved in acidified DMSO (5% 1N HCl in DMSO) to 5 mM. For animal experiments, VPC23109 was diluted into sterile normal saline (NS) and injected at 0.75 mg/kg.2

### Rodent studies

Mice were maintained on a 12:12-hr light-dark schedule in a temperature controlled specific pathogen-free facility. Diet induced obesity (DIO) mice (16-week old males) were purchased from Jackson laboratory (C57/BL6J DIO #380050, Bar Harbor, ME) and maintained on 60% kCal high-fat high-calorie diet (Research Diets, D12492, New Brunswick, NJ). Male mice were gavaged with either Dapa (1.25 mg/kg) or vehicle (Veh) for 4 days prior to LPS injection (10 mg/kg) at 8 pm on Day 4. *Apom*^*TG*^ and *Apom*^*KO*^ were previously described (9, 14). *Ly6g-cre LoxP-STOP-tdtomato* (Catchup mice), were previously described (24). Glucose was measured using a Glucocard Vital glucometer (Arkray USA, Minneapolis, MN). Creatinine and albumin were measured using the Liasys 330 (AMS diagnostics, Weston, FL). White blood cell and neutrophils were measured using a Hemavet 950 (Drew Scientific, Ramsey, MN).

### Cardiovascular Phenotyping

Transthoracic 2D imaging was performed in the Washington University Mouse Cardiovascular Phenotyping Core facility (https://mcpc.wustl.edu/) using a VisualSonics Vevo 2100 In Vivo Imaging System (Visual Sonics, Toronto, ON) according to the guidelines of the American Society of Echocardiography. Immediately prior to the procedure, mice were anesthetized with Avertin (2,2,2-tribromoethanol, 100 mg/kg, I.P., titrated to consciousness state and chosen due to its lack of cardio-depressive effects at the doses administered in this study). Echocardiography was performed on the morning after LPS administration, and subsequently read blindly by a cardiologist. Volumetric analyses were performed using the VevoStrain speckle-tracking software included in the VevoLab workstation (Fujifilm Visualsonics, Toronto, ON, USA) (25). For invasive hemodynamic studies, mice were anesthetized with Isoflurane (2% maintenance) + pancuronium (1 mg/kg given once). The mice were intubated and ventilated with a Harvard ventilator set at 200-400 μl. The right external jugular vein was cannulated with a 1.2 French high fidelity micromanometer pressure only catheter (SciSense Advantage System, Transonic, London, Ontario, Canada). The catheter was advanced into the right atrium to assess pressures. The ventilator was paused, and pressures were recorded and analyzed with SciSense analysis software in a blinded manner.

### Histologic analyses

For immunohistochemistry and tissue histology, kidney tissues were embedded immediately in cryomolds with optimal cutting temperature embedding medium, and frozen on dry ice. Immunohistochemical staining was performed with mouse anti-LRP2 (Abcam, ab76969, 1:200). Secondary antibodies were AF488 donkey anti-rabbit (Invitrogen A21206, 1:200, Carlsbad, CA, USA). Confocal images of fluorescent sections were obtained with a Zeiss Axio Imager M2 equipped with a Zeiss LSM 700 laser and a Fujitsu processor. Images were saved as CZVI files using Zen software (Black edition). Imagine analysis was performed using Visiomorph (VisioPharm, Broomfield, CO, USA). Quantification of mean fluorescence intensity was performed on 9 sections per mouse by NIH Image J software (http://rsb.info.nih.gov/ij/).

### Cell experiments

To produce ApoM-GFP, HepG2 cells were grown in 6-wells dish with DMEM and transfected using lipofectamine 3000 with a CMV driven construct containing the human *APOM* sequence and GFP. The expression of ApoM-GFP was verified by western blot analysis, microscopy, and qPCR. Cell medium was harvest after 48 hours and concentrated using Amicon ultra 30K spin columns. The concentration of apoM-GFP in collected cell medium was measured by EnSpire. HK2 cells were grown on 8-chamber slides for 24 hours. The cells were thereafter exposed to Dapaliflozin (6 nM) or vehicle for 16 hours, followed by addition of LPS (0.1 µg/ml) or saline as well as ApoM-GFP (2100 arb units) for further 6 hours. The cells were mounted with Fluoroshield. Images were obtained using a Zeiss Axio Scan.Z1. Quantifications of mean fluorescence intensity was performed by Image J software.

### Quantitative Real-Time Polymerase Chain Reaction analysis

Quantitative real time polymerase chain reaction (qPCR) was performed as described (25). Briefly, total RNA in kidney and liver tissues were extracted using RNeasy Mini kit (Qiagen, #74104, Hilden, Germany), and the first-strand cDNA was prepared using the iScriptTM cDNA synthesis kit (Bio-Rad, #1708890, Hercules, CA, USA). qPCR analysis was performed with SYBR Green Master Mix (Bio-Rad, #1725121) on QuantStudio 3 Real-Time PCR system (Applied Biosystems, A28136, Foster City, CA, USA) to determine relative mRNA levels. Mice sequences for qRT-PCR primers are shown below: ***Apom***: forward 5’-AAC AGA CCT GTT CTC CAG CTC G-3’, reverse 5’-GTC CAA GCA AGA GGT CAG AGA C-3’; ***S1pr1***: forward 5’-ACT ACA CAA CGG GAG CAA CAG-3’, reverse 5’-GAT GGA AAG CAG GAG CAG AG-3’; ***S1pr2***: forward 5’-CTC ACT GCT CAA TCC TGT CAT C-3’, reverse 5’-TTC ACA TTT TCC CTT CAG ACC-3’; ***S1pr3***: forward 5’-TTC CCG ACT GCT CTA CCA TC-3’, reverse 5’-CCA ACA GGC AAT GAA CAC AC-3’; ***Actb***: forward 5’-CAG AAG GAG ATC ACT GCC CT-3’, reverse 5’-AGT ACT TGC GCT CAG GAG GA-3’; ***36b4***: forward 5’-GCT TCG TGT TCA CCA AGG AGG A-3’, reverse 5’-GTC CTA GAC CAG TGT TCT GAG C-3’; ***Rpl32***: forward 5’-CCT CTG GTG AAG CCC AAG ATC-3’, reverse 5’-TCT GGG TTT CCG CCA GTT T-3’; ***Gapdh***: forward 5’-ACT CCC ACT CTT CCA CCT TC-3’, reverse 5’-TCT TGC TCA GTG TCC TTG C-3’.

### Western blot analysis

Western blot was performed on plasma, and tissue samples from mice as previously described (26). Primary antibodies employed were as follows: **mouse ApoM** (LS Bio, C158166, diluted 1:1000, Seattle, WA, USA), **LDLR** (Abcam, ab52818, diluted 1:500, Cambridge, MA, USA) and **β-Actin** (Sigma-Aldrich, A2066, diluted 1:4000, St. Louis, MO, USA). The band intensity was measured and analyzed with ImageJ software.

### Intravital Microscopy

*Ly6G-Cre LoxP-STOP-TdTomato* mice on chow diet were treated with vehicle or VPC23019 (0.75 mg/kg body weight (BW) IP daily) and gavaged with vehicle or dapagliflozin (1.25 mg/kg) for 4 days before IP injection of LPS (7.5 mg/kg). Twenty-four hours after LPS challenge, intravital microscopy was performed as we described (27). FITC-conjugated dextran (1 mg/g) was infused into a jugular venous cannulus. Neutrophil recruitment and vascular leak were monitored in an area of 0.02 mm^2^ (number/field/5 minutes) in the inflamed cremaster venules with a diameter of 25-40 μm. The number of neutrophils that visibly roll over the inflamed endothelium over 5 minutes were counted by an observer blinded to the treatment groups and unfamiliar with the study hypotheses. Adherent neutrophils were defined as neutrophils that were stationary for more than 30 seconds or crawled on the inflamed endothelium but did not roll over. Vascular leak was calculated by subtraction of the median fluorescence intensities of FITC-conjugated dextran inside of blood vessels from those of FITC-conjugated dextran in the field of view. The median extravascular FITC intensity per unit time (across the first 5 randomly selected vessels) and the per vessel area under the curve were calculated. In independent experiments, WT littermate controls, *Apom*^*KO*^, *Apom*^*TG*^ (along with their respective littermates) mice on chow diet were gavaged with vehicle or dapagliflozin for 4 days prior to LPS injection. Vascular leak and neutrophil recruitment were monitored by injection of FITC-conjugated dextran and Alexa Fluor 647-conjugated anti-Ly-6G antibodies (0.1 μg/g), respectively. Fluorescence and bright-field images were recorded using a Zeiss Axio examiner Z1 microscope system with a Yokogawa confocal spinning disk (CSU-W1) equipped with four stack laser system (405 nm, 488 nm, 561 nm, and 637 nm wave lengths). Images were collected with a high-speed, high-resolution camera (2304 × 2304 pixel format, ORCA-Fusion BT sCMOS, Hamamatsu). Data were analyzed using SlideBook, version 6.0 (Intelligent Imaging Innovations). Each curve is the median extravascular FITC-Dextran intensity across the quantified vessels (n = 5 vessels per mouse, 3-5 mice per group).

### Statistical analyses

Sample size was estimated based on literature review of murine SGLT2i studies (7, 28). Data were considered non-normally distributed if either the Shapiro-Wilk or Kolmogorov-Smirnov tests were statistically significant (*P* < 0.05). Data are presented as mean ± SEM or median ± 95% CI and analyzed by GraphPad Prism 9.0. Statistical significance among multiple groups was analyzed by analysis of variance (ANOVA) and multiple testing corrections as specified in each figure legend. In all of the analyses, a value of *P* < 0.05 was considered statistically significant.

### Study approval

All murine studies were approved by the Institutional Animal Care and Usage Committee at the Washington University in St. Louis and were performed following the Guide for the Care and Use of Laboratory Animals. Animal procedures were carried out in accordance with the Washington University School of Medicine Animal Studies Committee, which approved the protocols.

## Results

### Dapa preserves circulating protein levels of ApoM in LPS-treated mice by increasing proximal tubular Lrp2

To test our hypothesis that Dapa preserves circulating ApoM in acute inflammation, we utilized a model of diet-induced obesity (DIO) generated by feeding mice 60% high-fat high-calorie diet from 6 to 16 weeks of age. We performed studies in DIO mice, because it is a well-described model of pre-diabetes, obesity, and impaired glucose tolerance (29, 30), pathologies known to reduce ApoM (31, 32). DIO mice underwent oral gavage with vehicle or Dapa (1.25 mg/kg daily), prior to saline or LPS injection (10 mg/kg IP) on Day 4, followed by in vivo phenotyping (see below) and euthanasia on Day 5. LPS administration induced hypothermia, weight loss, and transient hypoglycemia, none of which was influenced by Dapa (**Supplementary Figure 1**).

In DIO mice, Dapa pre-treatment significantly attenuated LPS-induced decreases in circulating ApoM (**Figure 1A, B**). In contrast, Dapa did not have a significant effect on serum albumin concentrations (**Figure 1C**). We thus investigated how Dapa might attenuate LPS-induced reductions in ApoM. The liver, and to a lesser extent the kidney, are the main sites of ApoM production (18). We found that while LPS reduces both hepatic ApoM protein abundance and renal ApoM protein and mRNA abundance, none of these were affected by Dapa (**Figure 1D-I**). Since we did not find considerable change in ApoM mRNA and protein abundance in either the liver or the kidney, we sought to examine other potential mechanisms by which Dapa might preserve circulating ApoM.

**Figure 1.**
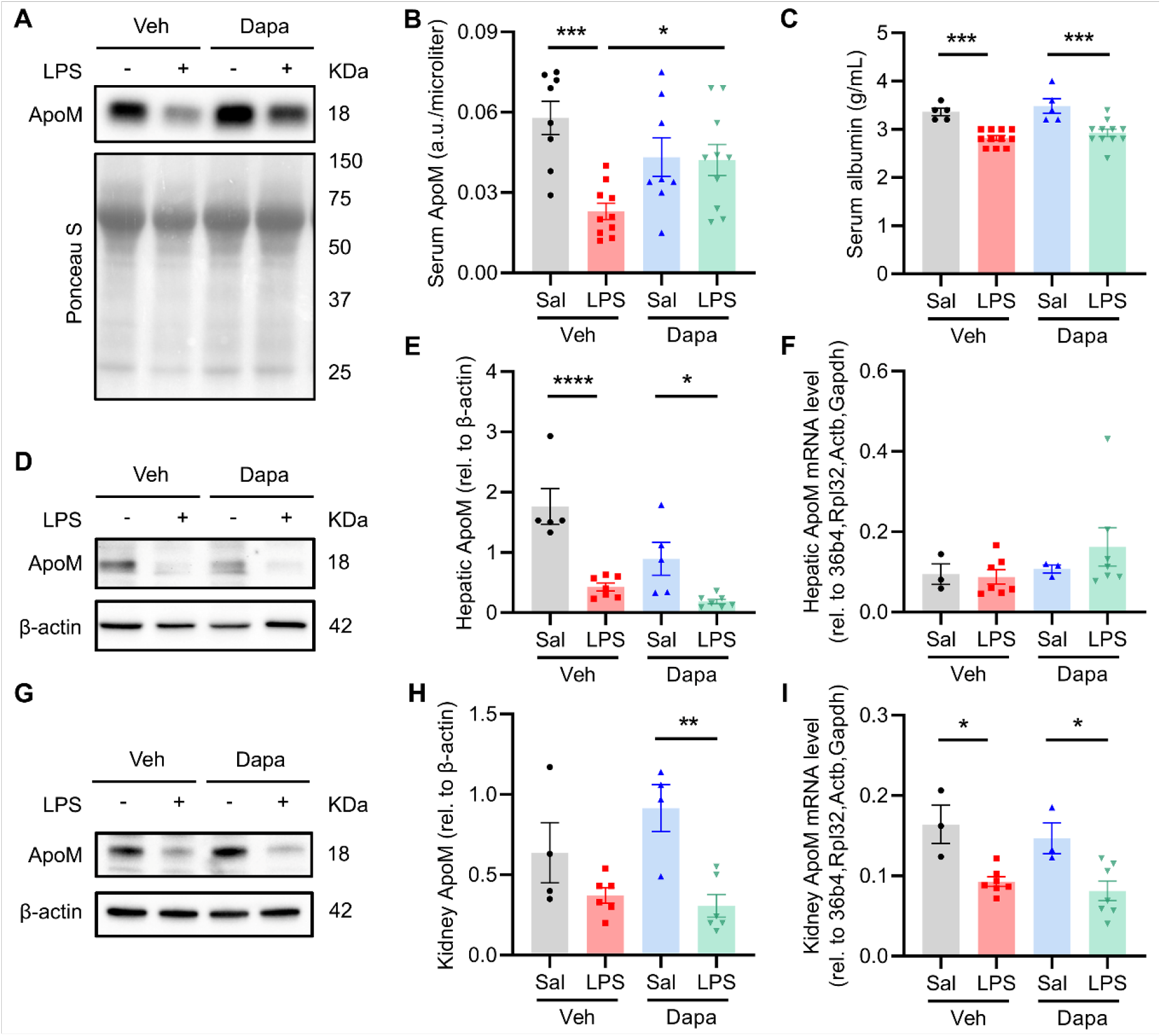
Dapagliflozin preserves circulating ApoM in LPS-treated mice. High-fat fed mice were gavaged with dapagliflozin or vehicle for 4 days before saline or LPS (10 mg/kg IP), and euthanized the next day, followed by: **A)** Western blot for murine ApoM from serum with quantification in **(B)** (n = 8-10); **C)** Serum albumin concentration (n = 5-11); **D)** Representative Western blot of hepatic ApoM with quantification in **(E)** and hepatic ApoM mRNA abundance in **(F)** (n = 3-7); **G)** Representative Western blot of kidney ApoM with quantification in **(H)** and renal ApoM mRNA abundance in **(I)** (n = 3-7). All data are presented as mean ± SEM. \**P* < 0.05, \*\**P* < 0.01, \*\*\**P* < 0.001, \*\*\*\**P* < 0.0001. Each dot represents one mouse. One-way ANOVA with Sidak’s correction for multiple comparisons in panels (B), (C), (E), (F), (H) and (I).

Hepatic ApoM clearance is dependent on low-density lipoprotein receptor (LDLR) uptake (33); however, LPS suppressed LDLR equally in both vehicle and Dapa-treated mice (**Figure 2A, B**), suggesting that hepatic ApoM clearance was unlikely to be responsible for the observed increase in circulating ApoM. Alternatively, we hypothesized that Dapa regulates ApoM reabsorption, which occurs in the renal proximal tubule by the protein receptor megalin (Lrp2) (34). Lrp2, a 600 kDa protein receptor and member of the low-density lipoprotein receptor family, is downregulated by high-glucose and LPS (35-37), and critical for ApoM reabsorption under conditions of kidney injury (38). Since the proximal tubule is also the site of SGLT2i/Dapa action, we hypothesized that Dapa may prevent LPS-induced reduction in Lrp2 levels. Blinded quantification of integrated fluorescent intensity from murine kidney sections stained with Lrp2 antibody showed that LPS-treated mice that received Dapa had higher levels of Lrp2 immunofluorescence compared to controls (**Figure 2C, D**). These findings suggest that Dapa can prevent LPS-induced reductions of Lrp2 in murine kidneys, potentially explaining how Dapa can preserve circulating ApoM in acute inflammation. This was further evaluated using human proximal kidney (HK2) cells treated with either LPS or LPS and Dapa. The uptake of ApoM-GFP was quantified and the results support that Dapa preserved the uptake of ApoM-GFP in renal proximal tubular cells under inflammatory conditions (**Figure 2E, F**).

**Figure 2.**
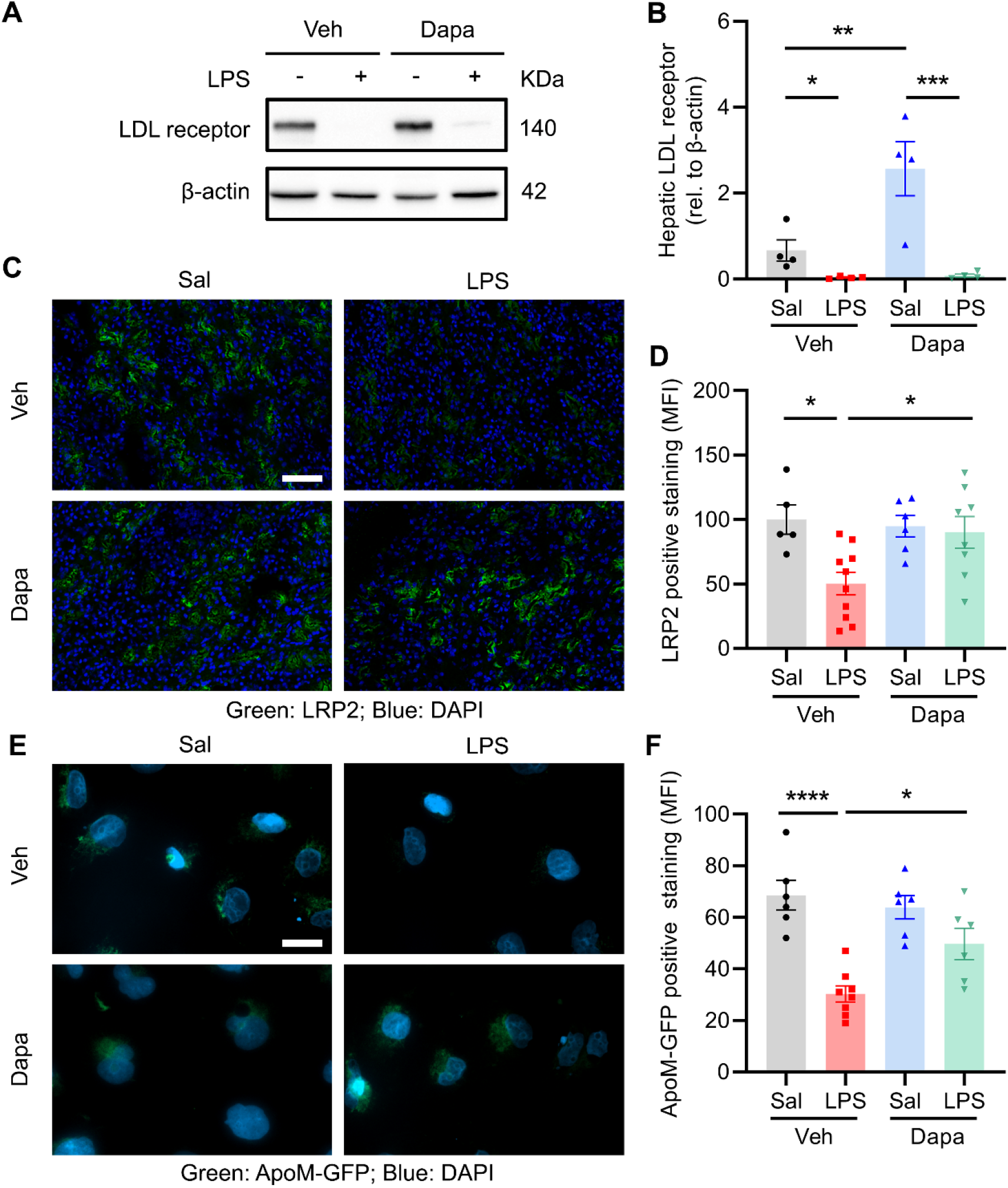
Dapagliflozin preserves circulating ApoM in LPS-treated mice by increasing proximal tubular Lrp2. High-fat fed mice were gavaged with dapagliflozin or vehicle for 4 days prior to LPS (10 mg/kg IP). **A)** Representative Western blot for LDL receptor and quantification in **(B)** (n = 4). **C)** Immunofluorescence staining for megalin/Lrp2 (Green) and DAPI in kidney frozen sections with quantification of mean fluorescence intensity in **(D)** (n = 5-11). Scale bar = 50 μm. **E)** Representative fluorescence images were obtained from HK2 cells which were grown on 8-chamber slides for 24 hours, thereafter, exposed to Dapaliflozin (6 nM) or vehicle for 16 hours, followed by addition of LPS (0.1 µg/ml) or saline as well as ApoM-GFP (2100 arb units) for further 6 hours. Scale bar = 20 μm. **F)** Quantification of mean fluorescence intensity in (E) (n = 6-8). All data are presented as mean ± SEM. \**P* < 0.05, \*\**P* < 0.01, \*\*\**P* < 0.001, \*\*\*\**P* < 0.0001. Each dot represents one mouse or one biological replicate (individual wells). One-way ANOVA with Sidak’s correction for multiple comparisons in panels (B), (D) and (F).

### Dapa attenuates cardiac dysfunction in LPS-treated mice via S1P receptor signaling

ApoM is inversely associated with survival and the acute phase response in HF patients (22), and improves end-organ function in LPS-treated mice (19). In our model, Dapa reduced serum creatinine (**Supplementary Figure 2**), consistent with prior data that SGLT2i attenuated renal injury in LPS-treated mice (7). Moreover, Dapa abrogated LPS-induced cardiac dysfunction (**Figure 3A**). Although LPS administration reduced heart rate in both vehicle and Dapa-treated mice (**Figure 3B**), echocardiography showed that Dapa attenuated LPS-induced reductions in ejection fraction, end-diastolic volume index, cardiac index, and coronary sinus area (**Figure 3C-F**). Invasive hemodynamic assessment of right atrial pressure showed that Dapa pre-treatment increased the pressure of atrial contraction, consistent with generally improved loading conditions and/or improved atrial function in LPS-treated mice (**Figure 3G, H**). Treatment of DIO mice with the S1P receptor 1 and 3 inhibitor VPC23019 (S1PRi) abrogated the effects of Dapa on cardiac ejection fraction, cardiac index, and coronary sinus area (**Figure 3I-K**), consistent with an important role for S1P receptor signaling in the effect of Dapa on cardiac function in LPS-treated mice.

**Figure 3.**
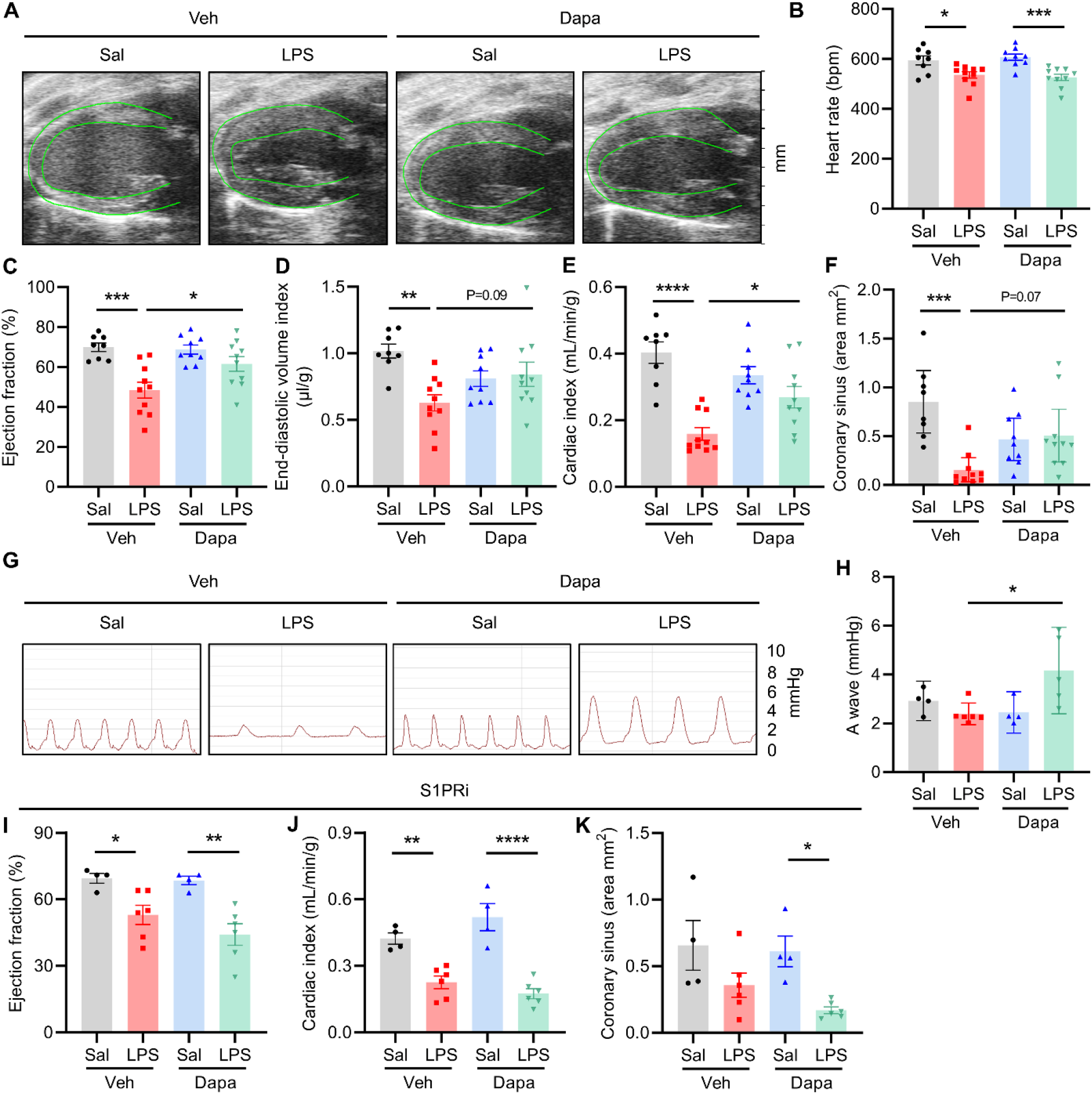
Dapagliflozin attenuates LPS-induced cardiac dysfunction in an S1P receptor dependent manner. High-fat fed mice were gavaged with dapagliflozin or vehicle for 4 days before saline or LPS (10 mg/kg IP) followed the next day by echocardiography under Avertin, titrated to consciousness. **A)** 2-D echocardiographic images, green line represents epi-and endocardial border as identified by speckle tracking, followed by blinded quantification of echo parameters (n= 8-10) including: **B)** Heart rate; **C)** Ejection fraction; **D)** End-diastolic volume indexed to body weight; **E)** Cardiac index; **F)** Coronary sinus area; **G)** Invasively determined right atrial pressure tracing with quantification in **(H)** (n = 4-6); **I-K)** High-fat fed mice were treated with the S1P receptor 1 and 3 inhibitor (S1PRi, 0.75 mg/kg IP daily), simultaneously gavaged with dapagliflozin or vehicle for 4 days before saline or LPS (10 mg/kg IP) followed the next day by echocardiography and quantitative assessment of ejection fraction, cardiac index, and coronary sinus area, respectively (n = 4-6). All data are presented as mean ± SEM or median ± 95% CI. \**P* < 0.05, \*\**P* < 0.01, \*\*\**P* < 0.001, \*\*\*\**P* < 0.0001. Each dot represents one mouse. One-way ANOVA with Sidak’s correction for multiple comparisons in panels (B-E) and (I-K), Kruskal-Wallis test with Dunn’s correction for multiple comparisons in panels (F) and (H).

### Dapa attenuates vascular leak and neutrophil transendothelial migration

One mechanism that exacerbates cardiac loading and function in sepsis is loss of endothelial barrier integrity and intravascular volume. We hypothesized that Dapa attenuates cardiac dysfunction in LPS-treated mice by reducing vascular inflammation and maintaining endothelial barrier integrity, thus preserving intravascular volume. To test this, we investigated the effects of Dapa on endothelial leak, as well as neutrophil rolling and adhesion on cremaster venules by intravital microscopy (IVM). We visualized the cremaster venules utilizing confocal IVM in chow fed *Ly6G-Cre LoxP-STOP-TdTomato* mice (24), in which neutrophils express a red fluorescent protein. Simultaneous injection of FITC-Dextran allows assessment of endothelial leak. Blinded quantification of fluorescent signal outside of the vessel showed that, compared with vehicle, Dapa treatment significantly decreased vascular leak (**Figure 4A, B** and **Supplementary Figure 3**). To examine whether the beneficial effect of Dapa is derived from S1P signaling, we tested the combined effect of Dapa and S1PRi. We found that compared with vehicle, S1PRi significantly increased vascular leak, and that treatment with S1PRi blocked the effect of Dapa on FITC-Dextran extravasation (**Figure 4A, B** and **Supplementary Figure 3**). Pre-treatment with Dapa increased the number of rolling neutrophils, and decreased the number of adherent neutrophils (**Figure 4C, D**); nonetheless, S1PRi blocked the effects of Dapa on rolling neutrophils, but not adherent neutrophils, implying that Dapa in part affects neutrophil recruitment in a S1PR-dependent manner. Importantly, we did not observe any consistent effects of Dapa on S1P receptor expression (**Supplementary Figure 4**), or significant increases in the number of circulating neutrophils (**Supplementary Figure 5**). These results indicate that Dapa prevents vascular leak induced by LPS, an effect dependent on S1PR signaling.

**Figure 4.**
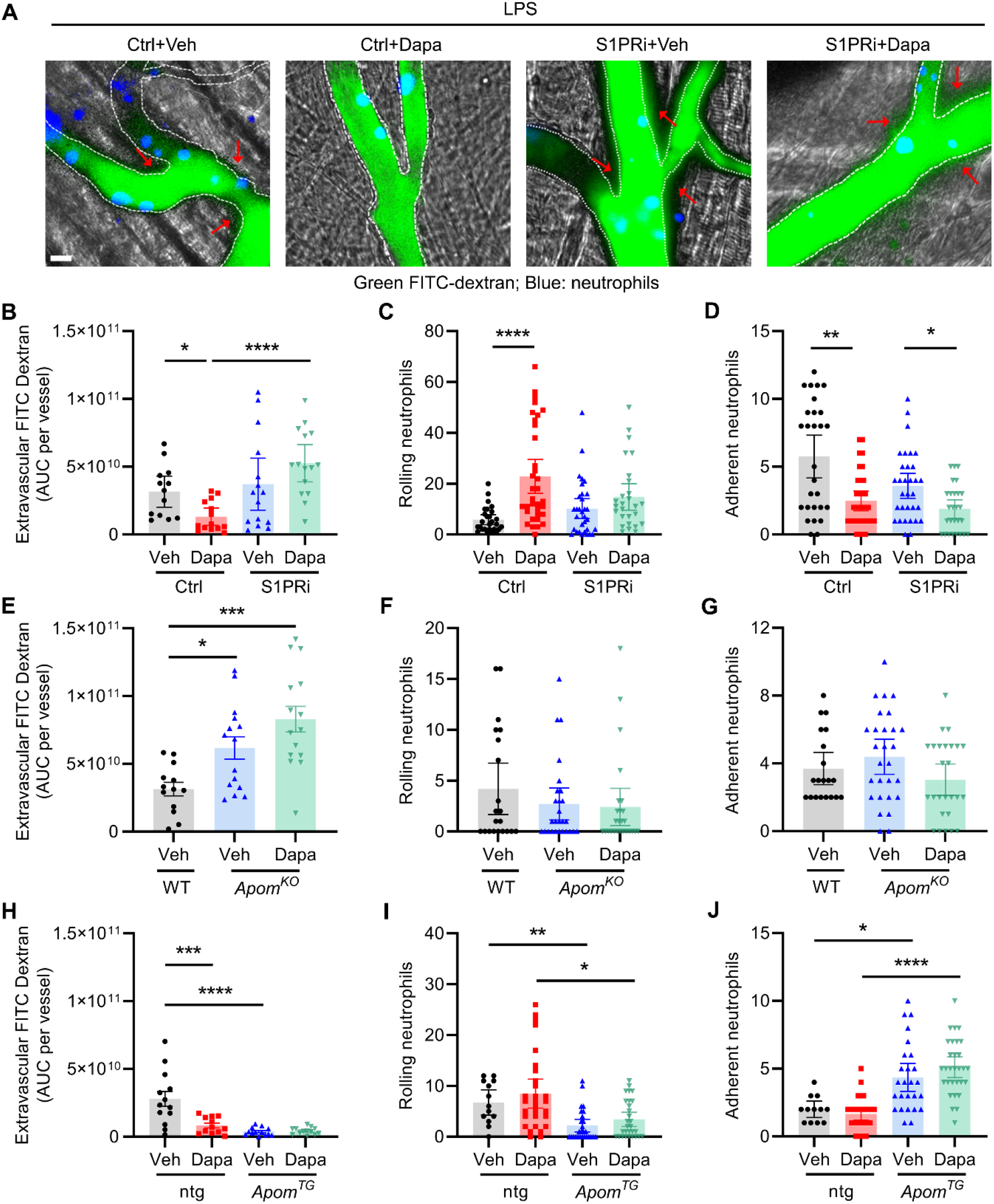
Dapagliflozin attenuates LPS-induced endothelial leak in an S1P and ApoM-dependent manner. *Ly6G-*Cre-TdTomato mice on chow diet were treated with control or S1PRi (0.75 mg/kg IP daily) and gavaged with dapagliflozin or vehicle for 4 days before LPS (7.5 mg/kg IP). Intravital microscopy was performed. **A)** Representative images of cremaster venules, FITC–Dextran (green), neutrophils (pseudocolor blue), white dotted lines indicate vessel border, and red arrows highlighting areas of extravascular FITC-Dextran, scale bar = 10 μm; **B)** Area under the curve (AUC) per vessel of extravascular FITC-Dextran; **C)** The numbers of rolling and **D)** adherent neutrophils observed in 5 minutes; **E-G)** Littermate control gavaged with vehicle for 4 days then treated with LPS compared to *Apom*^*KO*^ mice fed a chow diet and treated with vehicle or dapagliflozin gavage for 4 days prior to LPS injection. Intravital microscopy was performed to assess extravascular FITC-Dextran and rolling, and adherent neutrophils; **H-J)** Littermate control non-transgenic mice vs *Apom*^*TG*^ mice on chow diet were gavaged with vehicle or dapagliflozin for 4 days prior to LPS injection and intravital microscopy for assessment of extravascular FITC-Dextran and rolling and adherent neutrophils. All experiments quantified vessels (> 4 per mouse, n = 3-5 mice per group. Each dot represents one vessel. All data are presented as mean ± SEM or median ± 95% CI. \**P* < 0.05, \*\**P* < 0.01, \*\*\**P* < 0.001, \*\*\*\**P* < 0.0001. One way ANOVA with Sidak’s correction for multiple comparisons in panels (E) and (H), Kruskal-Wallis test with Dunn’s correction for multiple comparisons in panels (B-D), (F), (G), (I) and (J).

Given that Dapa prevented reductions in circulating ApoM in LPS-treated DIO mice, we sought to determine whether ApoM mediated the effects of Dapa on endothelial integrity and neutrophils. To test this, we utilized both ApoM knockout mice (*Apom*^*KO*^), which exhibit 50% reductions in plasma S1P (9), and mice with hepatocyte-specific overexpression of human *APOM* (*Apom*^*TG*^), which results in 3-5 fold increase in plasma ApoM and S1P (14). While *Apom*^*KO*^ mice exhibited increased vascular leak compared to littermate controls following LPS, Dapa did not significantly reduce FITC-Dextran extravasation in *Apom*^*KO*^ mice (**Figure 4E** and **Supplementary Figure 6**). Compared to control mice, *Apom*^*KO*^ mice did not affect neutrophil rolling and adhesion on the inflamed endothelium. Dapa pretreatment did not affect neutrophil rolling and adhesion in *Apom*^*KO*^ mice (**Figure 4F, G**). In contrast, IVM demonstrated that *Apom*^*TG*^ mice exhibit reduced vascular leak (**Figure 4H** and **Supplementary Figure 7**), fewer rolling neutrophils (**Figure 4I**), and more adherent neutrophils (**Figure 4J**), but Dapa pretreatment did not significantly alter any of these parameters in *Apom*^*TG*^ mice. These results suggest that Dapa reduces LPS-induced endothelial leak via Apom-S1P signaling, but that the effect of Dapa on neutrophil recruitment may only be partially dependent on ApoM/S1P.

## Discussion

We have uncovered that treatment with the SGLT2i Dapa attenuates cardiac dysfunction and endothelial leak, and preserves circulating ApoM in mice given LPS. We propose a model whereby, in the setting of systemic inflammation, Dapa preserves Lrp2 levels, leading to conservation of circulating ApoM, which in turn attenuates LPS-induced endothelial vascular permeability and endothelial inflammation. The most important finding of our study is that Dapa reduces LPS-induced endothelial leak in an ApoM/S1P dependent manner, which is of clinical importance in patients experiencing a severe inflammatory condition.

Our findings have multiple translational implications and link SGLT2i to the ApoM/S1P pathway, which is inversely associated with mortality in sepsis (10, 17-19), HF (22) and type 2 diabetes (10, 17-19, 23). ApoM also represents an important connection between diabetes and HFpEF, a condition where SGLT2i recently demonstrated benefits (39, 40). Clinical studies suggest that SGLT2i acutely reduce excess extravascular volume (41). We identify a mechanism by which SGLT2 inhibition in the proximal tubule preserves levels of ApoM, via increased re-uptake, contributing to preserved endothelial barrier integrity, therefore providing a potential explanation for how SGLT2i could acutely reduce excess extravascular volume compared to other diuretics. Finally, our results are of potential translational relevance given two ongoing trials of SGLT2i use in COVID-19 patients (NTC: NCT04381936 and NCT04505774), where ApoM is also inversely associated with clinical severity (20, 21).

One of the main findings of our study is that SGLT2i preserved cardiac preload, as evidenced by preserved end-diastolic volume index, coronary sinus area, and an increased right atrial contraction pressure during a severe inflammatory reaction. These hemodynamic effects are consistent with computer modeling and clinical observations that Dapa/SGLT2i reduce interstitial fluid, in particular in patients with fluid retention (42, 43). Moreover, it is known that ApoM reduces endothelial inflammation and preserves barrier integrity through endothelial S1P signaling (8, 9, 11, 44, 45). Results from our IVM studies show that extravascular leak was increased in ApoMKO but reduced in ApoMTG mice, with no additional effect of Dapa. Further, pharmacological inhibition of S1P receptors abrogated the effects of Dapa on endothelial barrier maintenance, demonstrating that the ApoM/S1P signaling pathway is necessary and sufficient for the effects of Dapa on endothelial leak.

Our results suggest that Dapa does not upregulate ApoM synthesis in the liver or kidney, the primary producers of ApoM. Rather, Dapa appears to preserve ApoM levels in the setting of an acute inflammatory reaction through reabsorption in the proximal tubule via upregulation of Lrp2. Although a prior study suggested that SGTL2i may reduce Lrp2 protein levels in a chronic type I diabetic mouse model (46), our present studies instead utilized both mice fed a high-fat high calorie diet or a chow diet. Interestingly, Lrp2 is downregulated both by hyperglycemia and LPS, conditions associated with urinary protein loss. Our finding that Dapa preserves Lrp2 is concordant with the clinical effects of SGLT2i, reducing renal protein loss in patients with type 2 diabetes (47). We posit that Dapa-mediated preservation of Lrp2 levels, in turn, protects against loss of ApoM, thus attenuating endothelial leak in acute inflammation.

Our work must be considered in the context of its limitations. First, we utilize an LPS model in both high-fat fed and chow-fed animals. We utilize lean animals in our IVM experiments in part because performing these studies in obese animals would add a significant technical challenge, and to avoid confounding from genetic interventions on the development of diet-induced obesity. While the LPS model itself is a generalizable model of cytokine storm, further studies in individual models of infectious versus sterile inflammation will be needed to extend our results to more specific clinical situations. In addition, our experimental design utilized a pretreatment strategy, which is relevant to patients either already taking a SGLT2i or to those who have recently been started on one. Further experiments will be needed to determine whether similar effects are observed if the SGLT2i is administered only after onset of severe inflammation.

In summary, our studies in Dapa treated mice given LPS show that Dapa a) preserves levels of ApoM, likely by preserving levels of Lrp2 in the proximal tubules; b) improves cardiac function in an S1P-receptor dependent manner; c) preserves endothelial barrier integrity via an ApoM/S1P mediated mechanism. These observations are translationally relevant and may stimulate future clinical studies of SGLT2i across a variety of disease states. Further translational and clinical studies will be required to determine whether Dapa or other SGLT2i prevent worse outcome in patients with sepsis of various etiologies, and whether our findings that Dapa preserves ApoM and endothelial barrier integrity play a pivotal role in chronic HF and kidney disease.

## Author contributions

Study conception: CVR, ZG, JO, RE, MK, AJ. Study design: CVR, ZG, TK, KNM, AK, CW, JO, RE, JS, MK, CC, JC, AJ. Data acquisition: CVR, ZG, TK, KNM, MO, AD, AG, LH, AK, CW, JN. Data interpretation: CVR, ZG, TK, KNM, MO, AD, AK, CW, JO, RE, MK, CC, JC, AJ. Manuscript writing: CVR, ZG, AJ. All authors accepted the manuscript.

